# Aging immune system-related blood cell DNA methylation can predict the onset of early stage colorectal cancer

**DOI:** 10.1101/2020.01.13.905299

**Authors:** Yuan Quan, Fengji Liang, Deqing Wu, Xueqing Yao, Yuexing Zhu, Ying Chen, Andong Wu, Danian Tang, Bingyang Huang, Ruifeng Xu, Zejian Lyu, Qian Yan, Zhengzhi Ning, Yong Li, Jianghui Xiong

**Author notes:** **Correspondence:** JianghuiXiong, Yong Li, Zhengzhi Ning, and Yuan Quan. These authors contributed equally to this work (Yuan Quan, Fengji Liang, Deqing Wu, Xueqing Yao).

## Abstract

**Objective:** There is a body of evidence that the aging immune system is linked to cancer. Here, we hypothesized that ubiquitination might play a key gatekeeper role in immune system aging and tumorigenesis in CRC. Therefore, we will systematically study the DNA methylation of ubiquitination genes and screen candidate CRC marker that are correlated with both aging and immune cell compositions.

**Design:** Through aging- and immune-related DNA methylation data, we investigated the DNA methylation regulation changes in promoters with other regions of genes during aging and their association with the immune cell proportion in the circulating whole blood of healthy individuals. Then, by collecting a cohort of 100 colon cancer patients and 50 healthy individuals, we used nucleic acid mass spectrometry to test whether ubiquitination genes can be used as candidate markers for the early screening of CRC.

**Results:** The biological analyses for aging- and CD4 T cell proportion-derived differential genes showed that they are associated with ubiquitination. Among them, DZIP3 was significantly associated with both aging (P-value = 3.86E-06) and CD4 T cell proportion (P-value = 1.97E-05) in circulating blood. Then, we validated that the 1^st^ exon DNA methylation of DZIP3 could predict the onset of early stage CRC (AUC = 0.833, OR = 8.82) and all pTNM stages of CRC (AUC = 0.782, OR = 5.70).

**Conclusions:** The epigenetically regulated ubiquitination plays an important role in immune aging and tumorigenesis. DNA methylation characteristic of DZIP3 can be used as a promising marker of CRC early screening.

**Summary box:** *What is already known about this subject?:*

1. The aging immune system is associated with cancer.
2. Ubiquitination plays potent roles in regulating a variety of signals in both innate and adaptive immune cells.
3. The abnormalities of ubiquitination are closely related to the occurrence of various tumors.
4. Blood cell DNA methylome analysis provides a promising tool to probe the key components of immune cell dysregulation in aging and tumorigenesis.

*What are the new findings?:*

1. The aging- and immune cell proportion-derived differential genes are associated with ubiquitination.
2. The epigenetically regulated ubiquitination plays an important role in immune aging and tumorigenesis.
3. DNA methylation characteristic of DZIP3, an E3 ubiquitin ligase with no reports on its function in immune cells and tumorigenesis, can be used as a promising markers of CRC early screening.

*How might it impact on clinical practice in the foreseeable future?:*

1. The in-depth study for DNA methylome of ubiquitination will open up a new path for the study of CRC pathogenesis, diagnosis and treatment.
2. In addition, DZIP3-focused screening technique only needs to extract the whole blood of individuals for MassARRAY analysis; the samples are easy to obtain, and the results are objective. Therefore, this method may have important applications in clinical CRC screening.

## 1. Introduction

In recent years, colorectal cancer (CRC) has become the third most common cancer in the world and is one of the leading causes of cancer-related death [1]. In the past few decades, due to a large number of studies on the pathogenesis of CRC, the diagnostic test and treatment strategies have made significant progress, which has led to the improvement of CRC survival [2].

Previous studies have found that there are several pathogenic factors of cancer, such as genome instability and mutation, intestinal flora imbalance and unreasonable diet [3]. Immune destruction is also one of the important causes of cancer. Therefore, chronic inflammation is one of the hallmarks of cancer [3], and CRC has long been known as one of the best examples of a tumor closely associated with chronic inflammation, which can be present from the earliest stages of tumor onset [4]. CRC arises following prolonged inflammation in the intestine, as in patients with inflammatory bowel disease (IBD), such as Crohn’s disease or ulcerative colitis [5]. Strikingly, CRC can be prevented or delayed by treatment with anti-inflammatory drugs [6, 7], which suggests the involvement of inflammatory processes in tumor onset. Inflammatory reactions are often linked to microbial responses, and the intestinal tract is populated by a myriad of bacterial strains that commonly live in harmony with their host, yet any substantial shift in the bacterial population can have a considerable effect on inflammatory responses and contribute to tumor development.

In addition, age is a major risk factor for many cancers. There is a body of evidence that aging immune system is linked to cancer [8–10]. There is a growing interest in immunosenescence and how it may contribute to the increased risk of cancer onset with aging. For example, CD4 T cells, especially T follicular helper cells, are critical for the generation of a robust humoral response to an infection or vaccination [11]. There is evidence that the aged microenvironment contributes to the age-related functional defects of CD4 T cells in mice [12], and the age-related impairment of the humoral response to influenza is associated with changes in antigen-specific T follicular helper cell responses [13].

Here, we hypothesize that ubiquitination might play a key gatekeeper role in immune system aging and tumorigenesis in CRC. Ubiquitination, the covalent attachment of ubiquitin molecules to proteins, is emerging as a widely utilized mechanism for rapidly regulating cell signaling. Recent studies indicate that ubiquitination plays potent roles in regulating a variety of signals in both innate and adaptive immune cells [14, 15]. A good example is that PD-1 ubiquitination could regulate the antitumor immunity of T cells. Dysfunctional T cells in the tumor microenvironment have an abnormally high expression of PD-1, and antibody inhibitors against PD-1 or its ligand (PD-L1) have become commonly used drugs to treat various types of cancer. A study showed that surface PD-1 undergoes internalization, subsequent ubiquitination and proteasome degradation in activated T cells [16].

The study of tumor immune interactions presents great technical challenges due to the great heterogeneity in the tumor microenvironment and global immune system function. Blood cell DNA methylome analysis provides a promising tool to probe the key components of immune cell dysregulation in aging and tumorigenesis. Several DNA methylation markers for the quantification of tumor immune cell infiltrates have been discovered, and their improved specificity over other markers has already been demonstrated. For example, DNA methylation of FOXP3 allows the distinction of T lymphocyte subpopulations such as regulatory T cells from other types of T lymphocytes. These subpopulations are difficult to distinguish by other methods (gene expression or immunohistochemistry) because they express the same surface markers [17, 18]. The power of DNA methylation as a tool for cell typing has further been underpinned by the development of DNA methylation signatures that allow the simultaneous quantification of different immune cell populations in blood. There are reports that a broad signature for hepatocellular carcinoma (HCC) in peripheral blood mononuclear cell and T cell DNA methylation, which discriminates early HCC stage from chronic hepatitis B and C and healthy controls, intensifies with the progression of HCC [19].

In the present study, we will systematically study the DNA methylation of ubiquitination regulatory system genes in blood cells and screen candidate genes that are correlated with both aging and immune cell compositions. Then, we will use low-cost nucleic acid mass spectrometry (Sequenom) to test whether ubiquitination genes can be used as candidate markers for the early screening of CRC.

## 2. Materials and Methods

### 2.1 Identification of differential genes associated with aging and immune cell proportions

In our study, aging-related (GSE40279) and immune-related (GSE69270) DNA methylation data were collected from the GEO database (www.ncbi.nlm.nih.gov/geo). GEO is an open-source functional genomics database of high-throughput resources, including microarray data, gene expression data, and DNA methylation data. The probes are transformed into the homologous gene symbol by means of the platform’s annotation information. The GSE40279 dataset contained DNA methylation profiles across approximately 450K CpGs in the human whole blood of 656 samples from patients whose age ranged from 19 to 101 years. The GSE69270 dataset contained DNA methylation profiles in the whole blood of 184 samples with annotations of six immune cell proportions, including CD8 T and CD4 T cells, monocytes, granulocytes, and NK and B cells. Next, we calculated the differential genes during aging and their association with immune cell proportion in the circulating whole blood of healthy individuals.

Recently, a study published in Genome Res. found that there is a strong relationship between the methylation of gene body difference to promoter (MeGDP) and gene expression, and the correlation coefficient is as high as 0.67 [20]. Therefore, MeGDP can be used as a predictor of gene expression. The higher the MeGDP is, the higher the expression value of a gene, suggesting that the degree of association with the relevant phenotype is higher. Based on the results of the above research, our other paper proposes the Statistical Difference of DNA Methylation between Promoter and other regions (SimPo) to calculate the DNA methylation characteristics of different genes (**Figure 1**) [21].

**Figure 1.**
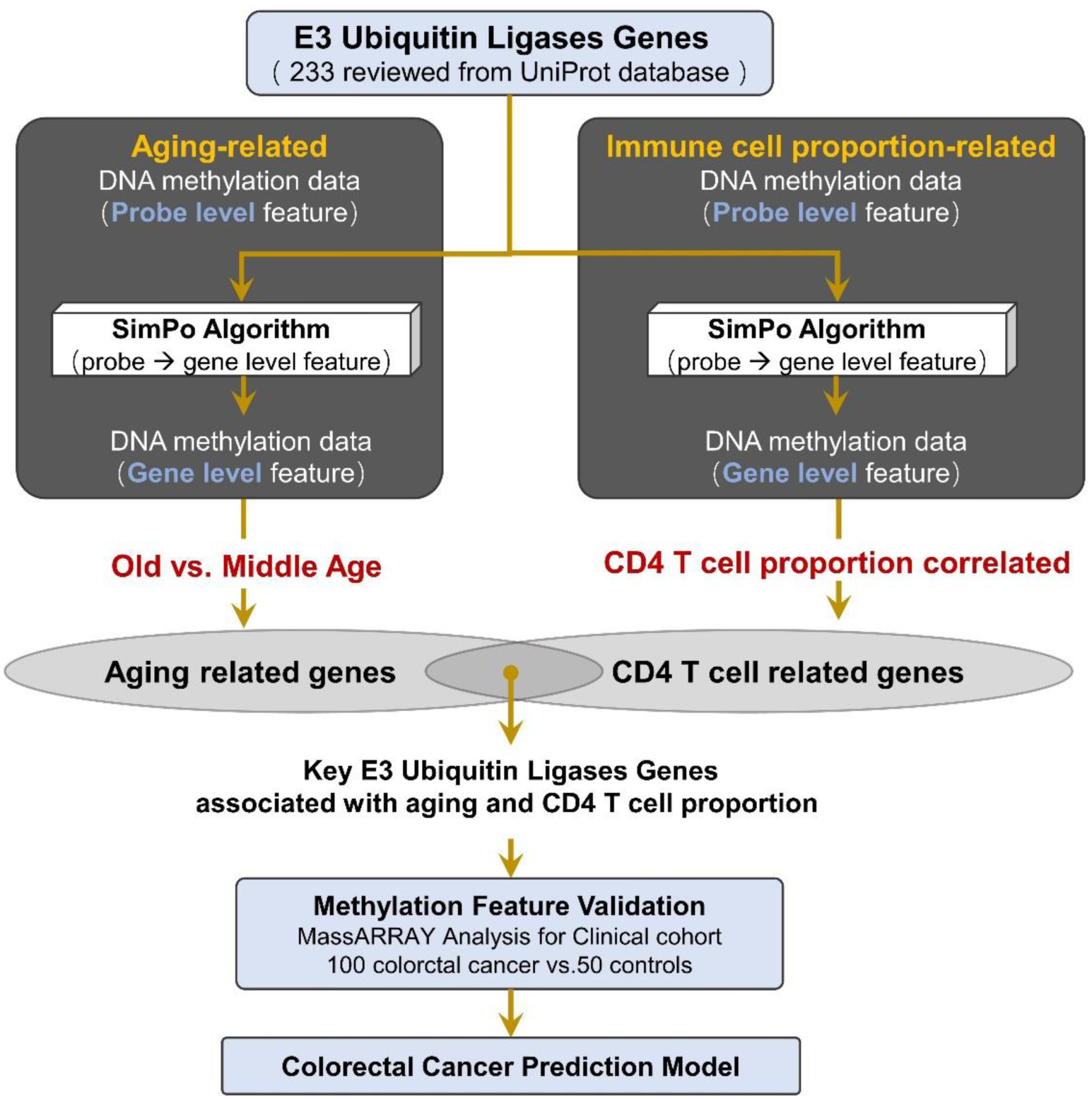
Method pipeline of candidate marker validation and prediction model construction for colorectal cancer.

The input data of the SimPo algorithm is the DNA methylation value of cg probes that are located in the gene promoter and the other regions. The significant difference method T-test is used in the SimPo algorithm, and the degree of difference (SimPo score) is used to characterize the DNA methylation of corresponding genes as follows [21]:

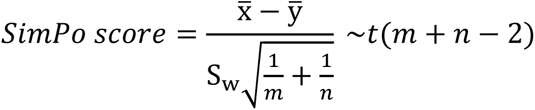

Where

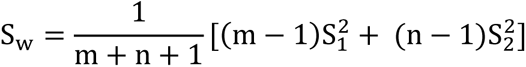

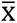: average DNA methylation value of all probes that are located in the promoter region; 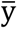: average DNA methylation value of all probes that are located in the other region; m: number of probes that are located in the promoter region; n: number of probes that are located in the other region; 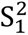: variance of DNA methylation values of probes that are located in the promoter region; and 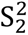: variance of DNA methylation values of probes that are located in the other region. In addition, since the SimPo score relates to the number of probes, to ensure the reliability of the SimPo score, we only selected genes with a number of promoter region-located and other region-located probes greater than or equal to five for further calculation.

Based on the SimPo algorithm, this study separately calculated the SimPo scores of each gene in the aging-related DNA methylation data and immune cell proportion-related DNA methylation data. Next, we calculated the differences in gene SimPo scores between the old age (> 50 year) and middle age (≤ 50 year) individuals based on the adjusted T-test. Then, for the immune-related methylation data, we identified the differential genes that were significantly associated with immune cell proportion based on the adjusted Spearman correlation test.

### 2.2 Ubiquitination genes collection

In this study, we collected 233 reviewed E3 ubiquitin ligase genes in humans by querying the UniProt database (https://www.uniprot.org) with “E3 ubiquitin-protein ligase” as a key word (**Figure 1, Table S1**).

The biological functions were enriched for 233 E3 ubiquitin ligases using the Kyoto Encyclopedia of Genes and Genomes (KEGG) pathway records in the Database for Annotation, Visualization and Integrated Discovery (DAVID, https://david.ncifcrf.gov/) [21]. The corresponding catalogs and diseases of the KEGG pathways were downloaded from the KEGG Pathway Database (https://www.genome.jp/kegg/pathway.html) [23]. The results showed that these E3 ubiquitin ligases were enriched in 4 KEGG pathways (Bonferroni-adjusted P-value ≤ 0.05), of which the most significant is the ubiquitin-mediated proteolysis pathway (**Table S2**). This result indicates that these 233 E3 ubiquitin ligases are significantly associated with ubiquitination and could be used for the next analysis.

### 2.3 Clinical blood samples collection

We analyzed blood samples from 100 diagnosed CRC patients (male: 67, female: 33) from Guangdong Provincial People’s Hospital and FoShan New RongQi Hospital. A total of 53% of the samples were diagnosed with pTNM stage I/II. We also collected 50 normal control samples (male: 18; female: 32) from FoShan New RongQi Hospital. All peripheral whole blood cell samples were treated with EDTA anticoagulant.

### 2.4 MassARRAY analysis

We identified the sequences 500 bp up and down from the CpG position at the 1st exon of DZIP3 (probe cg14787155 annotated position in HM450K chip) with the UCSC genome browser (http://genome.ucsc.edu/) (chr3:108308033-108309033), and designed 1 primer set for methylation analysis of the ***amplicon-cg14787155*** region by EpiDesigner software (http://epidesigner.com). Quantitative DNA methylation analysis of ***amplicon-cg14787155*** was carried out by using the MassARRAY platform (SEQUENOM) according to the official pipelines. In brief, DNA was treated with sodium bisulfite and PCR amplified, and bisulfite reactions were designed. The DNA methylation status of the samples was quantitatively tested by using the matrix-assisted laser desorption ionization time-of-flight mass spectrometry. Finally, 25 CpG sites were tested, and the methylation data of 18 individual units (one to two CpG sites per unit) were generated by the EpiTyper v1.0.5 software (Supplementary Material).

## 3. Results

### 3.1 Biological analysis of aging and CD4 T cell proportion-derived differential genes

Based on the SimPo algorithm, we calculated the SimPo scores for each gene of the aging and immune-related DNA methylation data. In the aging-related methylation data, we identified a total of 2358 genes that were significantly associated with the age of the patient (based on the adjusted T-test). In the immune-related methylation data, we identified a total of 1331 genes that were significantly associated with the CD4 T cell proportion (based on the adjusted Spearman correlation test).

First, we performed KEGG pathway enrichment analyses of the above significant gene sets by using the Enrichr database (https://amp.pharm.mssm.edu/Enrichr/) [24]. The results showed that the aging-associated genes were enriched in 32 significant KEGG pathways, and the immune-associated genes were enriched in 23 significant KEGG pathways. Among them, it is worth noting that the cellular senescence pathway is enriched in both the aging- and immune-associated gene sets (**Table 1**). In addition, the ubiquitin-mediated proteolysis pathway was identified by an immune-related gene set (**Table 1**). Therefore, the results indicated that genes that are significantly disturbed in DNA methylation level during the aging and immunization processes are associated with ubiquitination.

**Table 1.** The significantly enriched KEGG pathways of the aging- and CD4 T cell proportion-derived differential genes (top 20).

| Aging |  | CD4T cell proportion |  |
| --- | --- | --- | --- |
| KEGG Term | P-value | KEGG Term | P-value |
| Glyoxylate and dicarboxylate metabolism | 7.83E-05 | Ferroptosis | 1.04E-03 |
| Non-small cell lung cancer | 4.67E-04 | Cysteine and methionine metabolism | 3.40E-03 |
| Axon guidance | 1.11E-03 | Platelet activation | 8.16E-03 |
| Cell cycle | 2.39E-03 | **Cellular senescence** | **1.01E-02** |
| **Cellular senescence** | **6.67E-03** | Non-small cell lung cancer | 1.15E-02 |
| Nucleotide excision repair | 7.05E-03 | AMPK signaling pathway | 1.35E-02 |
| DNA replication | 7.18E-03 | TNF signaling pathway | 1.44E-02 |
| Valine, leucine and isoleucine degradation | 8.41E-03 | Hepatitis C | 1.50E-02 |
| Melanoma | 8.68E-03 | Hippo signaling pathway | 2.01E-02 |
| Human cytomegalovirus infection | 8.70E-03 | Base excision repair | 2.02E-02 |
| C-type lectin receptor signaling pathway | 9.24E-03 | Fatty acid degradation | 2.51E-02 |
| Dopaminergic synapse | 1.00E-02 | Viral carcinogenesis | 2.70E-02 |
| Oocyte meiosis | 1.07E-02 | Chronic myeloid leukemia | 2.87E-02 |
| Glioma | 1.29E-02 | Cushing syndrome | 2.91E-02 |
| Pancreatic cancer | 1.29E-02 | Tyrosine metabolism | 3.00E-02 |
| Hepatitis C | 1.40E-02 | Fatty acid elongation | 3.07E-02 |
| Chronic myeloid leukemia | 1.46E-02 | Progesterone-mediated oocyte maturation | 3.14E-02 |
| TNF signaling pathway | 1.72E-02 | Glycosaminoglycan degradation | 3.39E-02 |
| **Ubiquitin mediated proteolysis** | **1.72E-02** | Lysosome | 3.42E-02 |
| Hippo signaling pathway | 2.08E-02 | Leukocyte transendothelial migration | 3.51E-02 |

In addition, the Gene Ontology (GO) molecular function results further validate the above conclusion. The aging-derived differential genes were enriched in the ubiquitin-like protein ligase binding and ubiquitin protein ligase binding terms (**Table 2**). CD4 T cell proportion-derived differential genes were also enriched in the ubiquitin binding term (**Table 2**).

**Table 2.** The significantly enriched GO molecular function terms of the aging- and CD4 T cell proportion-derived differential genes (top 20).

| Aging |  | CD4T cell proportion |  |
| --- | --- | --- | --- |
| GO Term (Molecular Function) | P-value | GO Term (Molecular Function) | P-value |
| RNA binding | 8.10E-07 | Microtubule plus-end binding | 9.81E-05 |
| Protein-DNA loading atpase activity | 3.88E-04 | Kinase binding | 5.82E-04 |
| DNA clamp loader activity | 3.88E-04 | Phosphatidylinositol 3-kinase regulator activity | 1.10E-03 |
| Bubble DNA binding | 3.88E-04 | RNA binding | 1.70E-03 |
| Microtubule plus-end binding | 2.19E-03 | Protein kinase binding | 2.23E-03 |
| **Ubiquitin-like protein ligase binding** | **3.96E-03** | Clathrin adaptor activity | 2.96E-03 |
| **Ubiquitin protein ligase binding** | **4.26E-03** | Endocytic adaptor activity | 2.96E-03 |
| 5'-3' RNA polymerase activity | 7.76E-03 | Prenyltransferase activity | 4.41E-03 |
| Aminoacyl-trna ligase activity | 7.93E-03 | Transcription corepressor activity | 4.74E-03 |
| 3-hydroxyacyl-coa dehydrogenase activity (GO:0003857) | 9.11E-03 | Neurotrophin TRKA receptor binding | 5.05E-03 |
| Kinase activity (GO:0016301) | 1.32E-02 | RNA polymerase II transcription factor activity, sequence-specific transcription regulatory region DNA binding | 6.86E-03 |
| Protein kinase binding | 1.40E-02 | **Ubiquitin binding** | **8.33E-03** |
| Signal sequence binding | 1.55E-02 | Ribosomal protein S6 kinase activity | 8.40E-03 |
| Acetyltransferase activity | 2.21E-02 | Protein-DNA loading atpase activity | 8.40E-03 |
| Clathrin adaptor activity | 2.25E-02 | DNA clamp loader activity | 8.40E-03 |
| G-rich strand telomeric DNA binding | 2.25E-02 | Single-stranded DNA<br>endodeoxyribonuclease activity | 8.40E-03 |
| I-SMAD binding | 2.25E-02 | N-methyltransferase activity | 1.08E-02 |
| Magnesium ion transmembrane<br>transporter activity | 2.25E-02 | Hydrolase activity, hydrolyzing<br>N-glycosyl compounds | 1.14E-02 |
| Endocytic adaptor activity | 2.25E-02 | Ligand-dependent nuclear<br>receptor transcription coactivator<br>activity | 1.15E-02 |
| Double-stranded DNA binding | 2.37E-02 | Damaged DNA binding | 1.16E-02 |

The GO biological process results show that the aging-derived differential genes were enriched in the proteasome-mediated ubiquitin-dependent protein catabolic process and ubiquitin-dependent protein catabolic process terms (**Table 3**). The CD4 T cell proportion-derived differential genes were enriched in the positive regulation of the proteasomal ubiquitin-dependent protein catabolic process, the regulation of the proteasomal ubiquitin-dependent protein catabolic process and protein polyubiquitination (**Table 3**).

**Table 3.** The significantly enriched GO biological process terms of the aging- and CD4 T cell proportion-derived differential genes (top 20).

| Aging |  | CD4T cell proportion |  |
| --- | --- | --- | --- |
| GO Term (Biological Process) | P-value | GO Term (Biological Process) | P-value |
| DNA metabolic process | 3.23E-05 | **Positive regulation of proteasomal ubiquitin-dependent protein catabolic process** | **5.46E-05** |
| Cellular macromolecule biosynthetic process | 3.34e-05 | Positive regulation of proteasomal protein catabolic process | 3.28e-04 |
| Translation | 3.71e-05 | Cellular response to DNA damage stimulus | 3.55e-04 |
| Regulation of histone modification | 5.74e-05 | Negative regulation of protein serine/threonine kinase activity | 4.77e-04 |
| Cellular response to DNA damage stimulus | 7.21E-05 | Retinoic acid receptor signaling pathway | 5.81E-04 |
| Transcription, DNA-templated | 7.45E-05 | **Regulation of proteasomal ubiquitin-dependent protein catabolic process** | **7.40E-04** |
| Transcription-coupled nucleotide-excision repair | 8.76e-05 | Pyruvate metabolic process | 8.84e-04 |
| DNA repair | 1.18E-04 | Positive regulation of smooth muscle cell apoptotic process | 1.10E-03 |
| Nucleotide-excision repair | 1.25e-04 | Negative regulation of response to biotic stimulus | 1.48e-03 |
| **Proteasome-mediated ubiquitin-dependent protein catabolic process** | **1.56e-04** | Regulation of transcription from rna polymerase ii promoter | 1.61e-03 |
| Mitochondrial gene expression | 1.62e-04 | Dna repair | 1.73e-03 |
| Proteasomal protein catabolic process | 2.54e-04 | Negative regulation of protein phosphorylation | 1.84e-03 |
| RNA processing | 2.84E-04 | Toll-like receptor 3 signaling pathway | 1.88E-03 |
| Positive regulation of DNA biosynthetic process | 5.41E-04 | Negative regulation of cyclin-dependent protein kinase activity | 1.96E-03 |
| Gene expression | 6.40e-04 | Negative regulation of peptidyl-threonine phosphorylation | 2.21e-03 |
| TRNA aminoacylation for protein translation | 7.50E-04 | Regulation of defense response to virus by virus | 2.43E-03 |
| Positive regulation of chromatin silencing | 9.33e-04 | **Protein polyubiquitination** | **2.49e-03** |
| Regulation of endodeoxyribonuclease activity | 9.33e-04 | Hexose biosynthetic process | 2.50e-03 |
| Postreplication repair | 9.45e-04 | Response to laminar fluid shear stress | 2.96e-03 |
| **Ubiquitin-dependent protein catabolic process** | **1.03e-03** | G1 DNA damage checkpoint | 2.96e-03 |

Furthermore, we performed disease association analysis for the aging- and immune-related gene sets. Starting from the disease-gene associations of the ClinVar database [24], we found that the differential genes of aging were significantly associated with 12 diseases, and the differential genes of the immune system were significantly associated with 11 diseases. Interestingly, both are significantly associated with bowel cancer-related diseases, including colon carcinoma, hereditary nephrotic syndrome, and familial colorectal cancer (**Table 4**). These results indicated that aging and CD4 T cell proportion may play important roles in the development of colorectal cancer.

**Table 4.** The significantly enriched ClinVar diseases of the aging- and CD4 T cell proportion-derived differential genes (top 20).

| **Aging** |  | **CD4T cell proportion** |  |
| --- | --- | --- | --- |
| **Disease Term** | **P-value** | **Disease Term** | **P-value** |
| Hyperphosphatasia-intellectual disability syndrome | 2.37e-03 | Hyperphosphatasia-intellectual disability syndrome | 5.05e-03 |
| Diabetes mellitus type 2 | 3.68e-03 | Severe congenital neutropenia | 8.40e-03 |
| Familial hyperinsulinism | 9.11e-03 | Neoplasm of stomach | 1.28e-02 |
| Neoplasm of stomach | 9.11e-03 | **Familial colorectal cancer** | **2.31e-02** |
| Congenital central hypoventilation | 1.36e-02 | Combined oxidative phosphorylation deficiency | 3.39e-02 |
| **Carcinoma of colon** | **1.99e-02** | Adult junctional epidermolysis bullosa | 3.87e-02 |
| Beckwith-wiedemann syndrome | 2.49e-02 | Epidermolysis bullosa, junctional | 3.87e-02 |
| **Hereditary nephrotic syndrome** | **2.49e-02** | Familial adenomatous polyposis 1 | 3.87e-02 |
| Hirschsprung disease | 2.49e-02 | Galloway-mowat syndrome | 3.87e-02 |
| Combined oxidative phosphorylation deficiency | 2.51e-02 | Acute myeloid leukemia | 4.02e-02 |
| Neonatal diabetes mellitus | 3.97e-02 | **Carcinoma of colon** | **4.60e-02** |
| **Familial colorectal cancer** | **4.17e-02** |  |  |

### 3.2 E3 genes associated with aging

In the above results, we not only proved that the aging immune system is related to the onset of CRC but also further verified that ubiquitinated genes play an important role in the aging immune system. Therefore, we conducted further analyses on the 233 E3 genes. A total of 103 of the 233 E3 genes have cg probe annotations on the Illumina Infinium 450K Human Methylation Beadchip. Then, we obtained the SimPo scores of 42 E3 genes. The methylation SimPo scores of 23 E3 genes were significantly associated with the patient ages based on the adjusted T-test, and the E3 gene DZIP3 ranked sixth (**Figure 2a** and **Figure 3a**) (**Table S3**).

**Figure 2.**
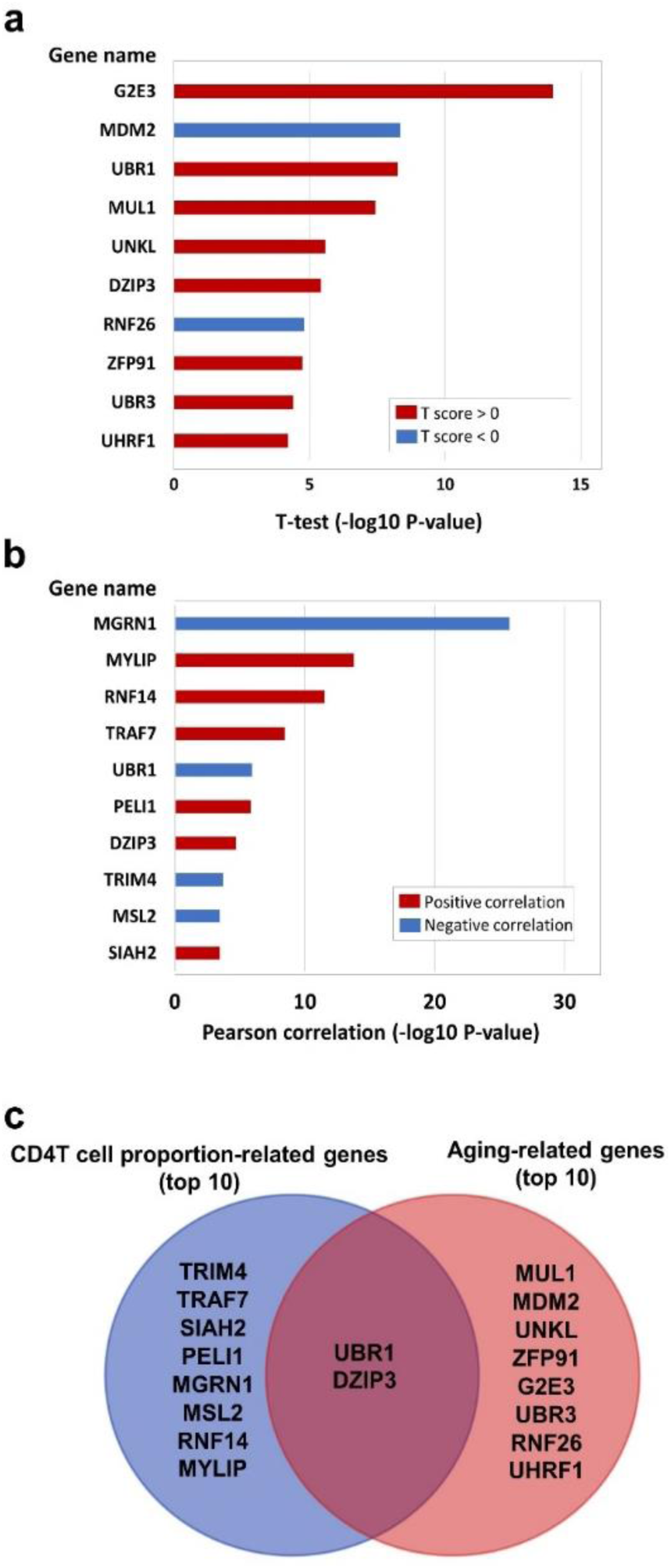
(a) The E3 gene list that is significantly associated with aging (top 10). (b) The E3 gene list that is significantly associated with CD4 T cell proportion (top 10). (c) The intersecting genes that are significantly associated with both aging (top 10) and CD4 T cell proportion (top 10).

**Figure 3.**
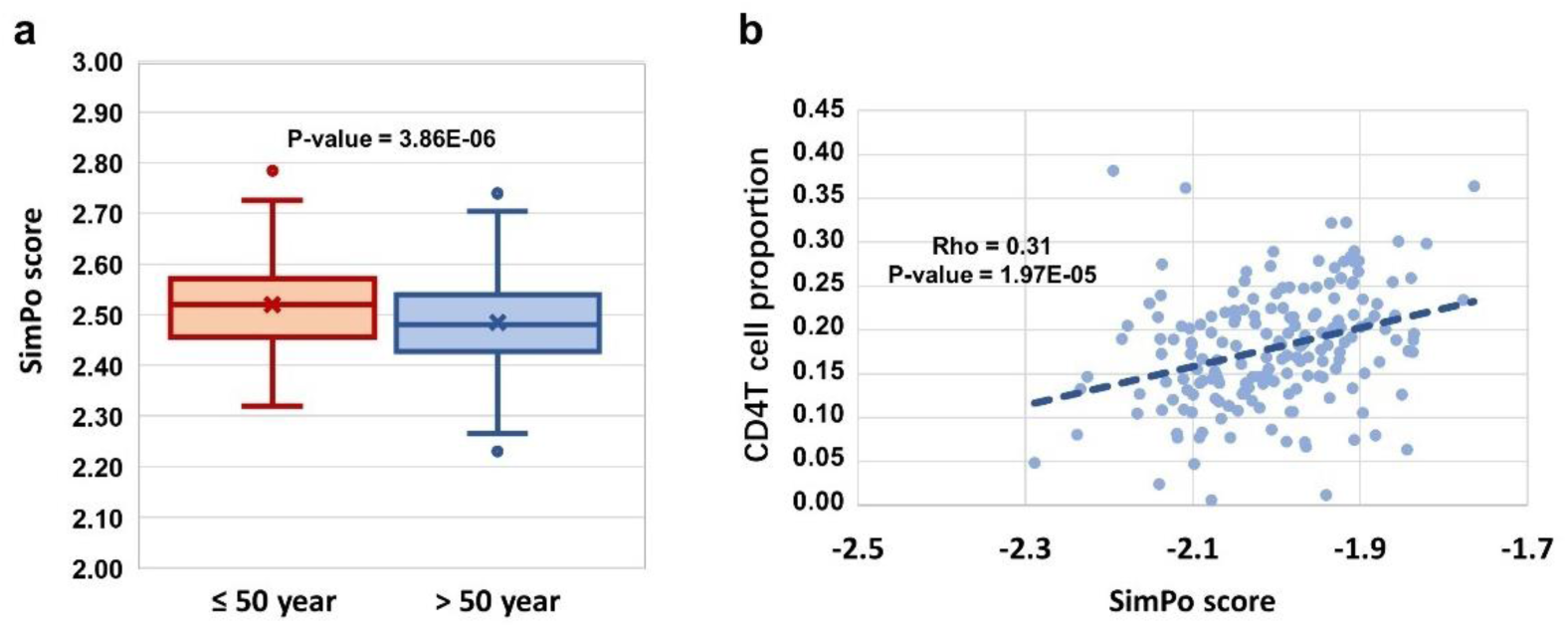
(a) DZIP3 is significantly associated with aging. (b) DZIP3 is significantly associated with the CD4 T cell proportion.

### 3.3 E3 genes associated with immune cell proportions

Then, we obtained the SimPo scores of 38 E3 genes based on the GSE69270 dataset. We calculated the Pearson correlations between the SimPo scores of the 38 E3 ubiquitin ligases with immune cell proportions (including CD8 T and CD4 T cells, monocytes, granulocytes, and NK and B cells) and the sum of CD8 T and CD4 T cells. The Pearson correlation results showed that 15 E3 genes were significantly associated with the CD4 T cell proportion (**Figure 2b** and **Table S4**), including the E3 gene DZIP3 (**Figure 3b**).

It is worth noting that there were two genes (UBR1 and DZIP3) that appeared in the top ten genes related to aging and CD4 T cell proportion (**Figure 2c**). Through literature searches in NCBI PubMed (https://www.ncbi.nlm.nih.gov/pubmed), it has been found that hundreds of published studies have demonstrated the associations of UBR1 with cancers. Interestingly, however, no article has reported the association of DZIP3 with colon cancer or even cancer. Considering the novelty of this gene, we conducted further clinical validation on DZIP3.

### 3.4 Methylation variations in DZIP3 amplicon-cg14787155

The clinicopathological characteristics of CRC patients are shown in **Table 5**. By MassARRAY analysis, we examined the methylation status of 25 CpG sites, and 15 methylation units were effectively detected in the majority of the samples for ***amplicon-cg14787155***. For each unit, we kept all the effective detected samples and tested the significance of methylation change in Ctrl vs. I&II and Ctrl vs. III. We found units showing significantly decreased methylation in colorectal cancer patients compared to normal controls (p < 0.05, T test). Specifically, we found that significantly changed methylation units were different between genders (**Figure 4A, Table S5**).

**Figure 4.**
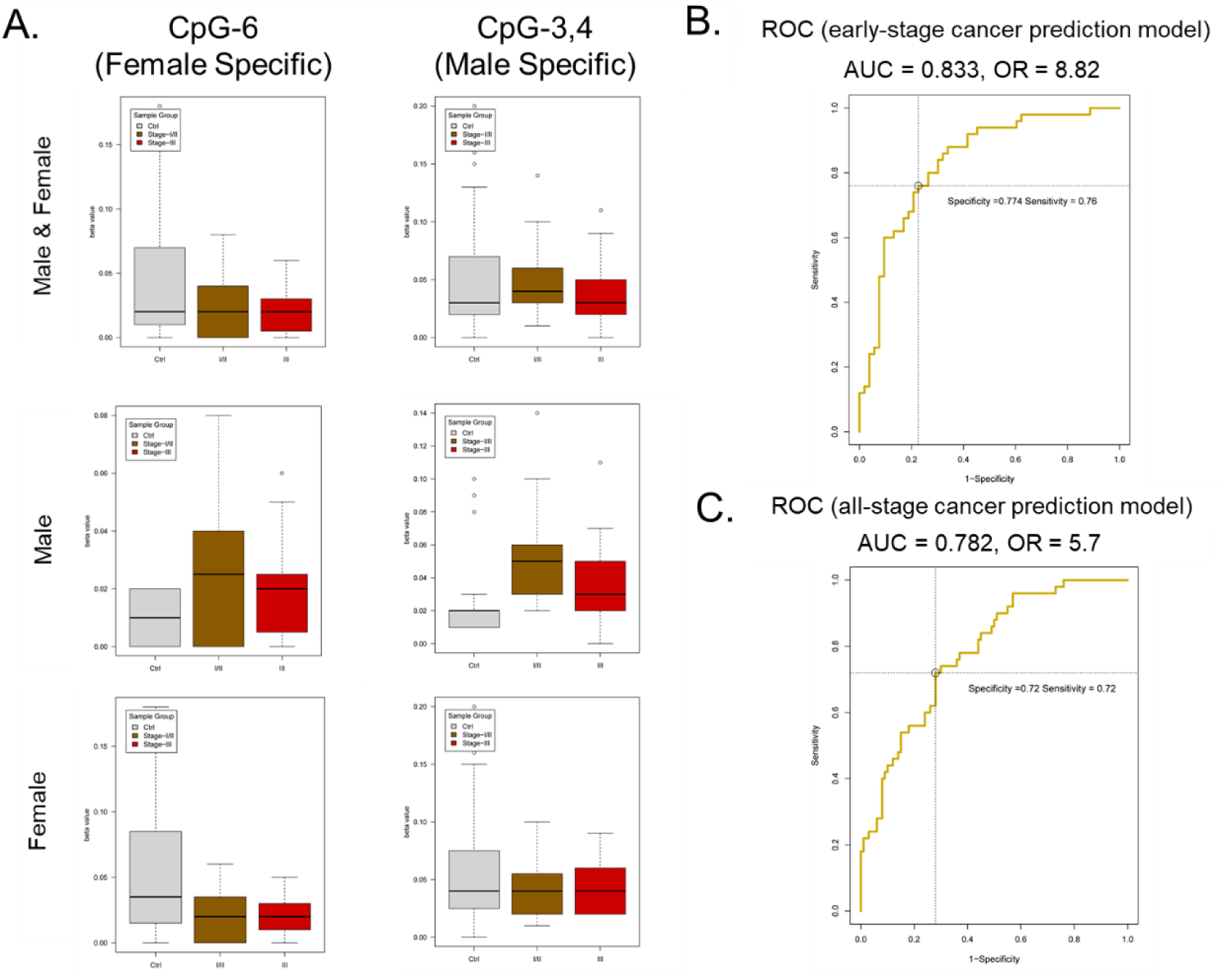
DNA methylation analysis of *amplicon-cg14787155* in DZIP3. (A) Sex-specific methylation level change between colorectal cancer and control in amplicon-cg14787155. (B) ROC analysis of the early-pTNM-stage colorectal cancer prediction model based on the MassARRAY data of ***amplicon-cg14787155*** features. The model with the best performance had an AUC of 0.833 and an OR of 8.82. (C) ROC analysis of the all-pTNM-stage colorectal cancer prediction model based on the MassARRAY data of ***amplicon-cg14787155*** features. The model with the best performance had an AUC of 0.782 and an OR of 5.7.

**Table 5.** Clinicopathological characteristics of the CRC patients (n = 150)

| Variable | Patients<br>(n = 100) | Control<br>(n = 50) |
| --- | --- | --- |
| Male/Female | 67/33 | 18/32 |
| Mean Age | 62.07 | 65.74 |
| Age range | 27 - 68 | 16 - 86 |
| TNM stage |  |  |
| I/II | 53 |  |
| III | 47 |  |

### 3.5 Colorectal cancer prediction model based on amplicon-cg14787155

With EpiDesigner software, we obtained the sequence of the MassARRAY primer relative to the amplicon-cg14787155 region (**Table 6**). Based on the effective detected methylation data of 15 methylation units, we constructed a prediction model to classify colorectal cancer patients from normal controls. As shown above, sex would be an important factor in predicting cancer state, and we also set sex as an input factor in the prediction model.

**Table 6.**
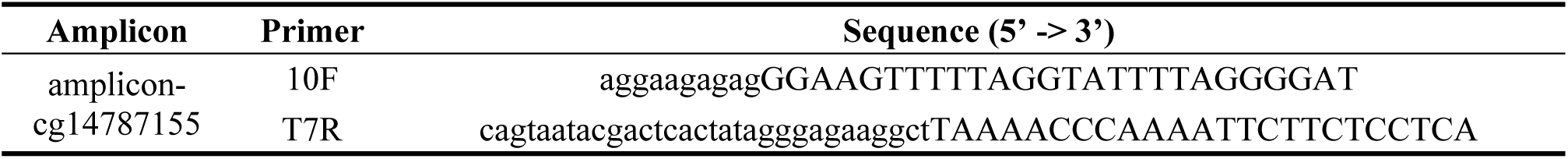
Sequence of MassARRAY primer and position relative to *amplicon-cg14787155*

A data set of 150 samples (100 colorectal cancer in stage I/II/III and 50 normal controls) with effective detected values in all 15 methylation units was kept for the model construction. The data were analyzed in R (v3.6) environment, the caret (v6.0-84) and ROCR (v1.0-7) packages were used to construct the logistic regression model, and leave-one-out cross-validation (LOOCV) was used to validate the model performance. First, we constructed the early-stage colorectal cancer prediction model in a dataset of 103 samples (53 colorectal cancer in pTNM stage I/II and 50 normal controls). The results showed that a model with the optimized input CpG sites (CpG_8, CpG_12, CpG_17, CpG_14.15, CpG_16, CpG_18, CpG_3.4, CpG_1.2, CpG_11, CpG_6) and sex showed the best performance (AUC = 0.833 and OR = 8.82) (**Figure 4B**).

Additionally, we constructed the all-stage colorectal cancer prediction model in the full data set. A model with the optimized input CpG sites (CpG_12, CpG_18, CpG_1.2, CpG_6, CpG_11, CpG_3.4) and sex showed the best performance (AUC = 0.782 and OR = 5.7) (**Figure 4C**). The results showed that the methylation state of the *amplicon-cg14787155* region has specific features in predicting early-stage and all-stage colorectal cancer.

## 4. Discussion

Ubiquitination is one of the main types of covalent modification of intracellular proteins. It participates in almost all life activities, such as cell survival, cell differentiation, and innate and acquired immunity [25–27]. The abnormalities of ubiquitination are closely related to the occurrence of various tumors [28–32]. It is worth noting that, consistent with the findings of this study, studies have shown that E3 ligase-mediated ubiquitination plays an important role in the development of CRC [33, 34]. The in-depth study of ubiquitination will open up a new path for the study of CRC pathogenesis, diagnosis and treatment. The results show that ubiquitination is an important regulatory mechanism of the surface proteins of immune cells and plays an important regulatory role in aging and changes in the composition of immune cells. The next step is to focus on whether DZIP3 plays an important role in the development of other types of cancer and on ubiquitinated pathway-modified proteins.

In this study, starting with E3 ubiquitin ligase genes, we screened early markers of CRC related to immune system aging and immune escape. The results showed that the ubiquitination regulatory gene DZIP3 was significantly correlated with age and changes in immune cell components. Furthermore, we designed a low-cost nucleic acid mass spectrometry kit, which was validated by recruiting patients with early colorectal cancer.

At present, the US FDA-approved CRC screening methods mainly include colonoscopy, fecal DNA testing, the fecal occult blood test (FOBT), the conventional blood carcinoembryonic antigen (CEA) indicator and the circulating methylated SEPT9 test. However, the above technologies still have certain limitations. For example, the early preparation of colonoscopy is cumbersome, invasive and subject to the subjective influence of the operator’s skills and clinical experience. Based on circulating methylated SEPT9 DNA for detecting colorectal cancer in a study by Church *et al.*, the methylation of SEPT9 DNA had a sensitivity of 35.0% for patients with stage I colorectal cancer, 63% for stage II, 46.0% for stage III and 77.4% for stage IV, with a specificity of 91.5% [35]. In recent years, peripheral blood circulating tumor cells (CTCs) have made great progress, and this technique is convenient for blood collection, with high public acceptance. However, there are still many problems in the application of CTCs. For example, positive results of CTCs still require confirmation by colonoscopy, and the appropriate testing interval of CTCs should be further explored.

However, this study used the DNA methylation characteristics of DZIP3 as early screening markers of early-stage CRC (sensitivity = 0.760, specificity = 0.774) and all-stage CRC (sensitivity = 0.720, specificity = 0.720). In addition, this screening technique only needs to extract the whole blood of individuals for MassARRAY analysis; the samples are easy to obtain, and the results are objective. Therefore, this method may have important applications in clinical CRC screening.

Another interesting feature of the present study suggests a novel early cancer screening approach based on blood cells. This blood cell methylation analysis mainly reflects the status of the immune system rather than cancer cells. Traditionally, liquid biopsy is focused on DNA mutation or DNA methylation tests on circulating free DNAs, which are mainly released by cancerous cells. Our results confirmed that aging and immune escape are important links in the development of intestinal cancer. Considering that a certain frequency of gene mutations will occur in healthy people and healthy tissues, the speed-limiting step of cancer occurrence, that is, the bottleneck, is not gene mutation but changes in the immune system. As cancer cells and immune systems reflect the yin-yang aspects of tumorigenesis, our results raise a new option for detecting the early and rate-limiting steps of tumorigenesis. Considering that blood cell methylation analysis could be conducted in healthy individuals, it is possible that our method could outline a continuous curve of healthy, inflammation, and transformation processes from enteritis to cancer. Thus, our method holds specific value for use in health management compared to the traditional liquid biopsy method, in which strong signals will appear only after enough cancer cells have been produced.

## 5. Ethics Statement

This study was approved by the institutional ethics committee of the Guangdong Provincial People’s Hospital and FoShan New RongQi Hospital (No.GDREC2016203H(R1)). The study was conducted in accordance with the ethical guidelines of the Declaration of Helsinki. Written informed consent was obtained from all participating patients before enrollment in the study.

## 7. Author Contributions

Yuan Quan and Fengji Liang conducted the data mining and bioinformatics analyses; Deqing Wu, Xueqing Yao, Zejian Lyu, Qian Yan performed the clinical sample and data collection; Yong Li and Zhengzhi Ning were responsible for the clinical trial design and organization; Zhengzhi Ning performed part of the clinical data mining; Yuexing Zhu, Ying Chen and Andong Wu took part in the lab experimental design; Danian Tang and Bingyang Huang took part in designing the clinical validation; Ruifeng Xu helped in preparing the manuscript; Jianghui Xiong led the epigenetic research and designed the strategy for the integrated analysis of DNA methylation data; Yuan Quan, Fengji Liang and Jianghui Xiong wrote the manuscript.

## 9. Availability of data and materials

The datasets supporting the results of this article are included within the article or in additional files. For the relevant raw data of clinical samples, we declare that the materials described in this manuscript will be freely available to any scientist wishing to use them for noncommercial purposes without breaching participant confidentiality.

